# Biomolecular condensation using *de novo* designed globular proteins

**DOI:** 10.64898/2025.12.19.695468

**Authors:** Andrey V. Romanyuk, T. P. Ragesh Kumar, Stephen J. Cross, Katarzyna Ożga, Rokas Petrenas, Joel J. Chubb, Jennifer J. McManus, Derek N. Woolfson

**Affiliations:** School of Chemistry, University of Bristol, Bristol, UK; Max Planck-Bristol Centre for Minimal Biology, University of Bristol, Bristol, UK; School of Physics, University of Bristol, Bristol, UK; Wolfson Bioimaging Facility, University of Bristol, Bristol, UK; NNF Center for Protein Design, University of Copenhagen, Department of Biology and Department of Drug Design and Pharmacology, Copenhagen, Denmark; Department of Chemistry, Wake Forest University, Winston-Salem NC, USA; School of Biochemistry, University of Bristol, Medical Sciences Building, Bristol, UK

## Abstract

De novo protein design is advancing rapidly, but many targets remain inaccessible to current AI-based tools. Here we describe *de novo* designed globular domains that drive biomolecular condensation. Starting from a water-soluble, monomeric protein, we make variants with the same amino-acid composition but different surface-charge distributions: one with large patches of surface charge, and another with a more-homogeneous charge distribution. The individual domains form stable and discrete structures in solution, with the large-patch variant exhibiting more-attractive interprotein interactions. Next, two copies of each variant are joined with disordered linkers to generate dumbbell-like proteins. When expressed in eukaryotic cells, the large-patch variant forms intracellular puncta, whereas that with small patches does not. The assemblies are dynamic, liquid condensates *in vitro* and in cells. The structured domains facilitate functionalisation: we introduce fluorophore-binding sites to visualise fluorescent condensates directly in cells without a GFP reporter; and we manipulate the condensates using motor proteins.

## INTRODUCTION

Despite major advances in *de novo* protein design^1,2^, several classes of target remain out of reach of current AI-guided methods. A prominent example is the complete de novo design of biomolecular condensates^3–8^, which requires control over weak multivalent interactions and the potential emergent phase behaviours. Existing protein-engineered condensates typically rely on natural scaffolds^9–17^, including intrinsically disordered regions^11,13,15,16^ or oligomerisation domains^14^. While fruitful, these strategies adapt rather than replace natural components. This leaves opportunities for developing truly bottom-up, fully designed, condensate-forming systems. If delivered, in principle, these would be amenable to rapid and predicable engineering to introduce functions to adapt them to different contexts.

Biomolecular condensates (BCs), also known as membrane-less organelles (MLOs), are subcellular compartments that play key roles in numerous biological processes, including: cell signalling, DNA organization, immune and stress response, metabolic reactions, and also in disease^18^. The biological roles of BCs stem from their hallmark dynamic behaviours, including small-scale motions^19^, molecular diffusion, component exchange with the surroundings^20^, and reversible assembly. Together with the lack of delimiting membranes, these properties enable rapid and adaptable responses to cellular conditions^18^. These unique features are highly attractive for synthetic biologists seeking new, robust and orthogonal approaches to generate functional microcompartments in artificial cells and engineered organisms, with eukaryotic cells presenting a particular challenge^21^.

BCs result from liquid-liquid phase separation (LLPS) of one or more macromolecular components, usually comprising proteins and/or nucleic acids. LLPS has been known to and understood by polymer and protein physicists for decades^22^. It occurs when intermolecular interactions are short-ranged and attractive^23^. However, its role in cellular biomolecular condensation has only been fully appreciated over the past two decades^24^.

From a structural perspective, intrinsically disordered regions (IDRs), globular domains, and linked-domain proteins can all undergo LLPS resulting in BCs in both cell-free systems and living cells^25^. Regardless of the protein organization, many of these systems have been described by sticker-and-spacer models, where stickers (e.g., short linear motifs, surface patches, or binding domains) drive attractive interactions with spacers separating these regions^25^. IDRs can be very efficient at condensation as they sample large conformational spaces giving residues high accessibility to make multivalent, dynamic interactions^26^. Globular proteins such as hemoglobin^27^, γ-crystallins^28^, and lysozyme^29^ can also undergo condensation in cells, though usually at higher concentrations^30^. Many BC-forming proteins (*e.g.,* hnRNPA1, FUS, LAF-1) have both IDRs and globular domains, with almost half of all proteins across the proteomes having such organization and even more in eukaryotes^31–33^. Phase separation of proteins with linked-domains or ‘beads-on-a-string’ structures with multiple folded sticker domains connected with flexible linkers (e.g. polySH3 in Nck, polySUMO, and polyubiquitin) play important roles in cell signaling^9^, protein homeostasis^34^ and autophagy^35^, respectively. Such proteins have inspired a range of synthetic linked-domain constructs of homo-^16^ and hetero-systems^9,14,36^ with multiple copies of natural oligomerizing or protein-binding domains. Nonetheless, the propensity of folded proteins to undergo LLPS in biological systems appears to be low.

Due to the above complexities and variations of natural BC-forming systems, the bottom-up engineering of phase-separating proteins is challenging^3^. However, there have been successes, for example: orthogonal translation enabled by spatial separation of components in synthetic organelles;^13,15^ and kinase-dependent formation of SPARK condensates^4^. The engineering/design considerations include: choosing the protein structure, *i.e.*, the use of IDRs or secondary and tertiary structures; selecting single or multi-domain organization; tuning the interplay between spacer and sticker modules; and directing the anisotropy of intra- and intermolecular interactions arising from the latter^23^. The problem is further complicated by the many factors that affect LLPS boundaries, including the properties of guest proteins that often need to be recruited for function. Until now, engineered BC-forming proteins largely rely on stickers from nature, e.g. from elastin^12^ or resilin^5^, from small linear motifs (SLiMs)^37^ to whole IDRs^11,13^ or folded domains^10^. This limits synthetic biology where building blocks orthogonal to those used by nature are preferable. An alternative to is to design proteins from scratch, i.e. completely *de novo*.

To date, *de novo* protein design has primarily focused on folded proteins with well-defined 3D structures. As a result, there are powerful approaches and tools for designing globular folded domains^2,38^. This has led to off-the-shelf building blocks that can be reprogrammed for function, including inducing protein-protein interactions (PPIs)^39,40^. Examples of *de novo* designed BCs are scarce. However, hybrid engineered-*de novo* systems^4,17,41^ and completely *de novo* phase-separating proteins (albeit containing conventional fluorescent proteins) derived from coiled-coil- or β-sheet-forming peptides have been reported^6–8^.

In contrast, here we combine completely *de novo* designed globular and IDR blocks to generate dumbbell-like proteins that undergo biomolecular condensation in eukaryotic cells, Fig. 1A. The globular domain is a hyperstable *de novo* 4-helix bundle (4HB). We use its surface as a canvass, adding small and large charged patches to programme weakly attractive but non-specific protein-protein interactions (PPIs). Two such domains are linked with *de novo* flexible linkers^7^ and a reporter fluorescent protein to give sticker-linker-reporter-linker-sticker constructs, Fig. 1A. The isolated domains are confirmed as stably folded, water-soluble proteins, with the large-patch variant having more-attractive interprotein interactions. Moreover, when duplicated in the dumbbell construct, only this variant leads to BCs in cells. In-cell fluorescence spectroscopy reveals dynamic and liquid-like condensates. Exploiting the designability of the system, we incorporate dye-binding sites into the 4HBs to allow direct visualization of fully *de novo* BCs in cells. Finally, we demonstrate that the system can be adapted to engage subcellular kinesin motors to transport the components and nucleate BCs at the cell periphery.

**Fig. 1:**
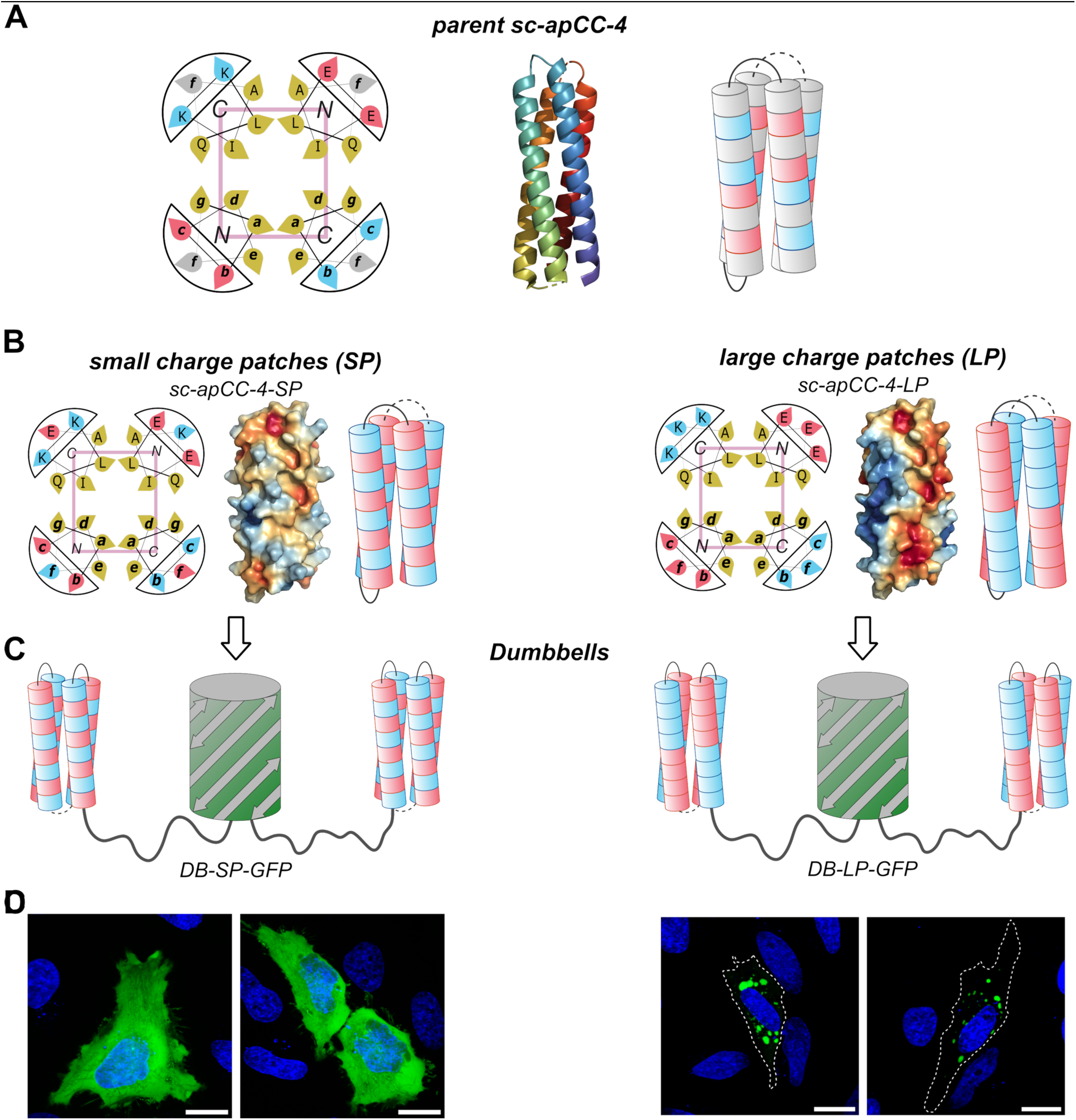
Rational design of 4-helix bundles and dumbbell proteins with small and large charged surface patches. A) Helical-wheel (left), X-ray crystal structure (middle, with the chain coloured chainbow from the *N* (blue) to *C* (red) terminus), and cartoon representations (right) of the parent sc-apCC-4 (PDB id, 8a3k). B) Helical-wheel (left), electrostatic potential surface maps generated with Adaptive Poisson-Boltzmann Solver (APBS) (middle, with the surfaces coloured from red (negatively charged) to blue (positively charged)), and cartoon representations of two variants with small (SP) and large (LP) charge patches. C) Dumbbell (DB) constructs with two copies of the designed modules fused to a fluorescent protein via 25-residue long disordered linkers. D) Live confocal fluorescence images of HeLa cells transiently expressing DB-SP-GFP (left) and DB-LP-GFP (right).

## RESULTS AND DISCUSSION

### *De novo* design of globular domains for biomolecular condensation

Net charge^42^, charge blockiness in IDRs^43^, patchiness of globular-protein surfaces^36,44–46^, and charged-residue content^47^ are all known to influence protein solubility and diffusion^42^, PPIs^43^, and protein-polyelectrolyte interactions^48^. Therefore, we tested if electrostatic interactions alone could be used to design a *de novo* dumbbell BC-forming system.

To generate a folded multivalent module as a sticker, we started with a *de novo* designed, single-chain, antiparallel 4-helix bundle, sc-apCC-4 (Fig 1A; PDB ID 8a3k), built using well-understood sequence-to-structure relationships^49^. The helices have heptad sequence repeats, (*gabcdef)_4_*. The *g, a, d* and *e* sites define the core helix-helix interactions and were maintained in the design. The *b, c* and *f* positions are on the surface and are generally mutable^50^. We exploited this through rational design to generate eight variants, each with 24 negatively charged glutamic acid (Glu, E) and 24 positively charged lysine (Lys, K) residues but distributed differently to give various charged patches (Fig. 1B & Extended Data Fig. 1, and Supplementary Table 1).

Next, we combined two copies of each of the eight patchy variants in a “sticker-spacer-sticker” manner to form dumbbell-like (DB) proteins (Fig. 1C & Extended Data Fig. 1C). The spacer comprised two copies of a flexible 25-residue peptide^7^ flanking the mEmerald variant of green fluorescent protein (GFP) as a reporter. We screened these constructs for phase separation directly in eukaryotic cells (Fig. 1D & Extended Data Fig. 1C). The designed proteins showed dramatically different behaviours when transiently expressed in HeLa cells. Live-cell imaging using confocal fluorescence microscopy of DB proteins carrying relatively small patches (designs 1 – 4, Extended Data Fig. 1A&B) revealed diffuse fluorescence throughout the cell except the nucleus (Extended Data Fig. 1C). By contrast, designs with larger patches (designs 5 – 7) formed distinct puncta around the nuclei, and the variant with the largest charged patches (design 8) showed extremely small speckles with substantial amounts of the protein present in the nuclei.

For detailed characterization, we focused on two variants with clearly different and representative phenotypes: from the diffuse set, we chose design 1, which had a relatively homogeneous charge distribution (hereafter called small patches, “SP”); and from the set that formed puncta, design 6, which had an anisotropic charge distribution (large patches, “LP”) (Fig 1B&D, Extended Data Fig. 1). The dumbbell constructs for these were named DB-SP-GFP and DB-LP-GFP (Fig. 1C). The HeLa phenotypes of these were maintained in HEK cells (Extended Data Fig. 2), where the puncta formed by DB-LP-GFP were particularly large (up to ≈10 μm) (Extended Data Fig. 2B). Moreover, the DB-LP-GFP puncta behaved as liquids exhibiting coalescence, fission, and flow in live cells (Extended Data Fig. 2C, Supplementary Video 1). To test for any subcellular localization of DB-LP-GFP, live-cell imaging with organelle staining was used. This showed that the puncta did not colocalise with any particular compartment, *e.g.* mitochondria, lysosomes, ER, actin filaments, or nucleus (Supplementary Fig. 1). Finally, to test that the sticker-spacer-sticker construct was critical, three different fusions of a single LP domain with GFP were transiently expressed in HeLa cells (Supplementary Fig. 2). Unlike the DB construct, DB-LP-GFP, none of these formed puncta indicative of bimolecular condensates.

### Large charged patches drive liquid-like biomolecular condensates

The individual patchy domains (SP and LP) were expressed in *E coli* and purified using immobilised metal affinity chromatography (IMAC) and size-exclusion chromatography SEC (see Methods). The *N*-terminal histidine tags were removed to eliminate any contributions to PPIs. Despite the large number of changes made from the parent protein (16 point mutations each for the SP and LP designs) and the exclusively Glu/Lys-based surfaces, solution-phase characterization of both variants confirmed stable, discrete and fully folded proteins consistent with the designs: The circular dichroism (CD) spectra were typical of largely α−helical proteins (Fig. 2A), and variable-temperature CD measurements indicated highly thermostable folds (Supplementary Fig. 3). Analytical size-exclusion chromatography (SEC) gave sharp single peaks indicating monodisperse proteins (Supplementary Fig. 4), and analytical ultracentrifugation (AUC) returned molecular weights of 15.3 kDa (0.98 × monomer) and 16.3 kDa (1.04 × monomer) for the SP and LP proteins, respectively (Supplementary Fig.5). We could not crystallise either design. However, AlphaFold2 and AlphaFold3 predictions (Supplementary Table 2) had confident metrics and largely matched the X-ray crystal structure of the parent sc-apCC-4 protein. Furthermore, the AlphaFold models for the SP and LP proteins fitted small-angle X-ray scattering (SAXS) data with χ^2^ values of 1.07 and 1.03, respectively (Fig. 2B, Supplementary Table 2).

**Fig. 2.**
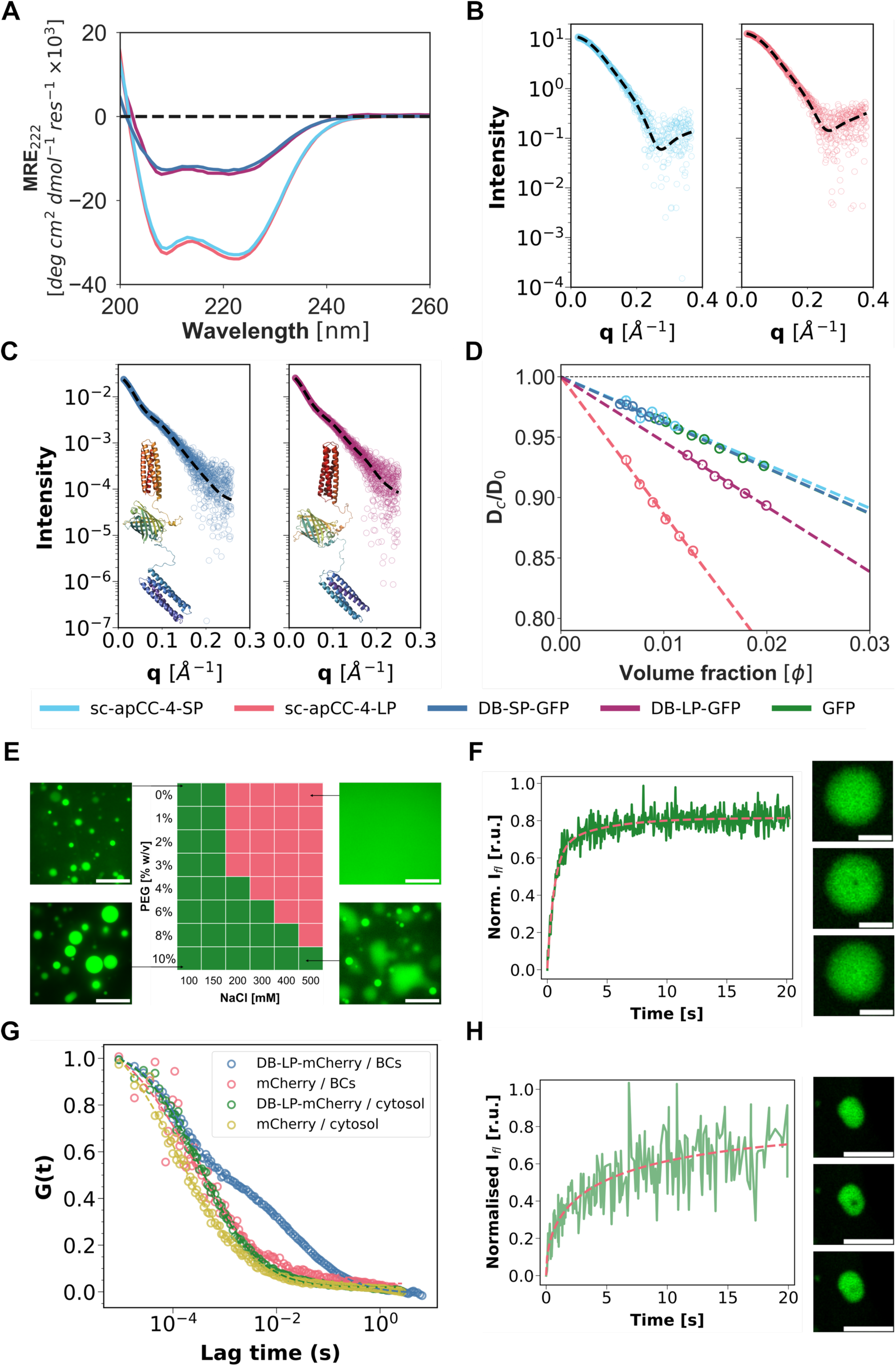
Characterization of designed patchy sc-apCC-4 variants and dumbbell constructs. A) CD spectra of sc-apCC-4-SP and sc-apCC-4-LP (both at 25 µM), and DB-SP-GFP and DB-LP-GFP (both at 5 µM) at 5 °C. B) SAXS data for sc-apCC-4-SP (left, χ^2^ 1.07) and sc-apCC-4-LP (right, χ2 1.03) at 1 mg/mL protein fitted to their AlphaFold2 models using FoXS. C) SEC-SAXS data for DB-SP-GFP (left, χ^2^ 1.01) and DB-LP-GFP (right, χ^2^ 0.97) at 5 mg/mL protein fitted to their AlphaFold2 models with allowed flexible linkers using MultiFoXS. The best-fit models are presented under the plots. D) The dependence of *D_C_/D_0_* (*D_C_*, collective diffusion coefficient; *D_0_*, free-particle diffusion coefficient) on protein volume fraction for SP, LP, DB-SP-GFP, DB-LP-GFP, and GFP measured by DLS at 20 °C. Colour key for (D-G): SP, cyan; LP, red; DB-SP-GFP, blue; DB-LP-GFP, purple; and GFP, green. Conditions: 50 mM Tris pH 7.5, 500 mM NaCl. E) Screen of NaCl and PEG concentrations that result in either single-phase (red) or two-phase liquid-liquid systems (green) for DB-LP-GFP in 50 mM Tris-HCl solution at pH 7.5. Fluorescence images show examples of de-mixed liquid droplets and a homogeneous solution. Scalebars = 5 μm. F) FRAP in the de-mixed droplets formed by DB-LP-GFP protein (4 mg/mL) in 50 mM Tris-HCl pH 7.5, 150 mM NaCl, and 2% PEG3350 at 37 °C. The average data and fitting to a bi-exponential function are in green and red, respectively. Representative images of a protein droplet before and at 40 ms and 5 s after bleaching. Scalebars = 5 μm. G) The FCS diffusion data for the fluorescent probes (DB-LP-mCherry or mCherry) in either DB-LP-GFP droplets or cytoplasm presented as the normalised autocorrelation curve. H) FRAP in BCs formed by DB-LP-GFP in live HeLa cells. The averaged data and fit to a biexponential function are in green and red, respectively. Representative images of a protein droplet in cells before and at 40 ms and 20 s after bleaching. Scalebars = 5 μm. All experiments performed with HeLa cells were at 37 °C.

DB-SP-GFP and DB-LP-GFP were also expressed from *E. coli*, purified, and characterised as described for the isolated SP and LP domains. Again, the CD spectra were typical of α−helical proteins (Fig. 2A), with MRE values consistent with ≈50% of the protein being intrinsically disordered or β sheet from the linker and GFP domains, respectively, and the proteins had high thermostabilities (Supplementary Fig. 3C&D & 6). SEC and AUC measurements for DB-LP-GFP confirmed a monodisperse species with a molecular weight of 64.7 kDa (1.02 × monomer) (Supplementary Fig. 4, 5C & 7). SAXS data for DB-SP-GFP and DB-LP-GFP fitted well to their AlphaFold2 predicted structural models (Fig. 2C, Supplementary Table 2). Moreover, MultiFoXS fitting to these data gave models with radii of gyration of 4.80 and 4.75 nm, respectively, consistent with the hydrodynamic radii of 4.5 – 5.0 nm measured by dynamic light scattering (DLS; Fig. 2C and Supplementary Fig. 8; see Methods).

Next, to probe the potential for interprotein interactions in the individual SP and LP domains, we measured their diffusion coefficients using DLS at different protein concentrations to determine the net interaction parameter (diffusivity, *k_D_*), (Fig. 2D)^51^. Negative values of *k_D_* indicate net attractive PPIs. These were more attractive for the anisotropic variant, LP, with *k_D_* = -11.4 ± 0.5 (pink line) compared with *k_D_* = -3.4 ± 1.2 (blue line) for SP. Extending this to the DB constructs, for DB-LP-GFP *k_D_* = -5.4 ± 0.2 (Fig. 2D). Thus, its interprotein interactions are less attractive than for its individual 4HB domain—reflecting the addition of GFP and the IDR linkers—but this value is similar to natural proteins that undergo LLPS^28^. For DB-SP-GFP *k_D_* = -3.8 ± 0.3, similar to its component domains, indicating that intermolecular interactions are unlikely to be sufficiently attractive to produce LLPS.

To probe the potential for the SP and LP domains to form specific PPIs, we turned to MaSIF (molecular surface interaction fingerprinting – see Methods), which assesses chemical and geometric features on protein surfaces. Despite the different charge arrangements, neither variant was predicted to have hot spots indicative of tight PPIs (Supplementary Fig. 9). Also, using PROPKA, the pK_a_ of the multiple acidic and basic ionizable side-chain groups in both variants were predicted to be at least two units below and above pH 7.5, respectively, resulting in the nearly identical predicted net charges of - 2.32 and -2.28 (Supplementary Fig. 10). Therefore, the differences in *k_D_* reflect the spatial arrangements of surface residues rather than from a difference in the net charge.

On this basis, the LP variant appears to be a good candidate for a *de novo* globular domain with non-specific attractive interactions typical of those associated with dynamic liquid-like behaviour in biomolecular condensates^52,53^.

### Dumbbell proteins form liquid-like, dynamic biomolecular condensates

To explore condensate formation by DB-LP-GFP *in vitro*, we screened for solution conditions that produced liquid-liquid phase separation (Fig. 2E, Extended Data Fig. 3). The pH was fixed at 7.5 with Tris-HCl buffer, and NaCl and PEG (MW 3350) concentrations were varied to probe the effect of ionic strength and molecular crowding, respectively. At up to 150 mM NaCl, in the absence of a crowding agent, we observed liquid–liquid de-mixing with characteristic formation of spherical macroscopic droplets. This seems counter-intuitive given the net negative charge and relatively low ionic strength. We posit that electrostatic attraction arises from the anisotropic charge distribution designed into this protein, as observed for engineered natural proteins^46^. At higher ionic strengths, the protein charge is screened, reducing the contribution of electrostatic attraction to the interprotein interactions, since LLPS is only observed with added PEG, producing a depletion attraction. Surprisingly, over the range of ionic strengths and PEG concentrations screened, we did not observe any other condensate types (e.g. aggregation), suggesting that there may be a subtle interplay between electrostatic attraction and electrostatic repulsion driving phase separation for this charge variant.

Next, we probed the dynamics of the de-mixed DB-LP-GFP droplets *in vitro* using fluorescence recovery after photobleaching (FRAP) in buffer with 150 mM NaCl and 2% PEG (Fig. 2F). The data fitted best to a bi-exponential function, with an overall *τ*_1/2_ = 0.53 s. This is significantly lower than half-times (of several to tens of seconds) observed for de-mixed droplets formed by FUS^54^, DDX4 *N-*terminal IDR^55^, LAF-1^56^, hnRNPA1^57^, and NPM1^58^ (Supplementary Table 3), and indicates highly mobile proteins in our *de novo* designed system (Supplementary Fig. 11, Supplementary Table 4). To test for similar liquid-like behaviour in cells, we measured FRAP in live HeLa cells 24 hours after transfection with DB-LP-GFP DNA (Fig. 2H). The observed recovery was almost as complete as *in vitro*, with a slightly longer effective *τ*_1/2_ of ≈1.2 s that was best fit using either bi-exponential or anomalous diffusion models (Supplementary Fig. 12, Supplementary Table 5). We attribute the difference in the viscoelastic behaviour between a solution and the cell microenvironment to the high sensitivity of the phase boundary to the salt concentration (Fig. 2E, Extended Data Fig. 3) due to the large charged patches on protein surface. There is also a possibility of some maturation within the droplet over the 24-hour period between cell transfection and imaging.

To investigate the dynamics within the droplets in cells further, we used fluorescence correlation spectroscopy (FCS) to measure diffusion based on fluorescence fluctuations of an mCherry fluorescent probe. This was introduced either in the dumbbell variant, DB-LP-mCherry, or as free mCherry co-expressed alongside DB-LP-GFP (Fig. 2G). To compare diffusion in the dense phase of the DB-LP-GFP condensates and a dilute phase (i.e., proteins outside of droplets), fluorescence fluctuations of the probe were recorded in droplets in cells, and in the cytoplasm of cells from the same experiment with an mCherry signal but without obvious condensates. This gave autocorrelation curves (Fig. 2G) from which diffusion coefficients (Supplementary Table 6) were measured by fitting to an anomalous diffusion model (Fig. 2G, dashed lines).

Diffusion of the mCherry-only construct both inside (droplets) and outside (cytoplasm) the condensates indicated a single mobile fraction in each case, but with different diffusion coefficients of 34.3 ± 7 and 66.5 ± 6 µm^2^/s, respectively. Thus, as expected, the probe diffused more slowly in the condensates. Moreover, the value for the free, cytoplasmic mCherry was consistent with those reported for EGFP in the cytoplasm of HeLa cells at 50 – 56 µm^2^/s (Supplementary Table 3). Similarly, the non-condensed cytoplasmic DB-LP-mCherry behaved as single mobile fraction, albeit with slower diffusion coefficient (23.7 ± 0.9 µm^2^/s) than the free mCherry probe, as expected for a larger protein.

By contrast, within the BCs, the diffusion of DB-LP-mCherry showed fast-moving (72.1 ± 2.6 µm^2^/s) and slow-moving (0.36 ± 0.02 µm^2^/s) mobile fractions (Supplementary Table 6). The fast component is consistent with the diffusion of monomer in a viscoelastic background (like the measurement of monomer in the cytoplasm). Assuming a monomeric DB-LP-mCherry probe, we estimated the viscosity of the condensates to be ≈ 143 ± 9 mPa·S (Table S6). This is comparable to and within the liquid-like, dynamic regime observed for FUS^54^, DDX4^55^, LAF-1^56^, and NPM1^58^ (Table S3). The slow component indicates compositional heterogeneity within the BC^59^. In contrast, free mCherry showed single-component diffusion, excluding a gel-like droplet state. These observations support reversible association of the DB-LP-GFP protein within condensates, but not in the dilute phase, consistent with short-range attractive interactions and with our design hypothesis: compared to free mCherry, diffusion of DB-LP-mCherry in condensates is slowed by the sticky LP domains introduced to drive attractive interactions and BC formation.

### Dumbbell condensates can be functionalised for direct imaging in cells

An advantage of phase-separating systems made from folded *de novo* domains rather than IDRs is that the former can be more readily functionalised. Recently, we developed a computational design pipeline for endowing *de novo* protein modules with small-molecule-binding sites^60^. This gave a variant of sc-apCC-4 that binds the cell-permeable fluorophores Nile Red (NR) and Nile Blue (NB), sc-apCC-4-NR-1, with binding constants of K_D,NR_ = 2.40 ± 0.06 μM and K_D,NB_ = 0.59 ± 0.28 μM, respectively. Based on this, we replaced hydrophobic-core residues in the first LP domain of DB-LP-GFP with the binding-site residues from sc-apCC-4-NR-1 to give DB-LP-GFP-NR-1, Fig. 3A. Thus, the first module is bi-functionalised: on its surface for driving phase separation, and in its core for fluorophore binding.

**Fig. 3.**
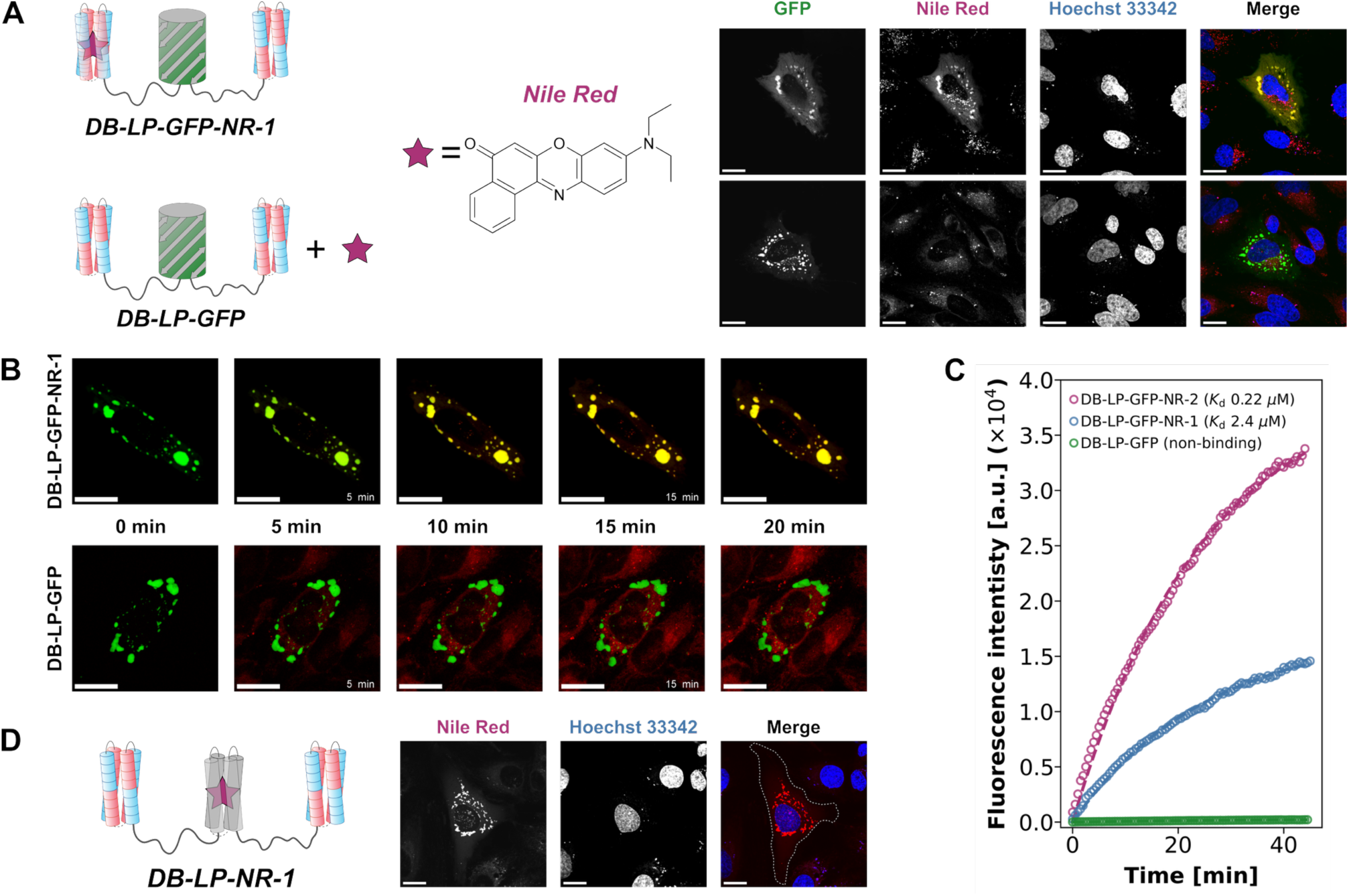
Confocal fluorescence images of live HeLa cells with BCs functioning as “molecular sponges” for Nile red (NR). A) DB-LP-GFP-NR-1 contains a bifunctional *de novo* domain for LLPS and NR binding. Its BCs are stained by NR (row 1). BCs of the non-NR-binding parent, DB-LP-GFP, do not uptake NR (row 2). B) Time-lapse imaging of NR binding by DB-LP-GFP-NR-1 compared to the non-binding control DB-LP-GFP. C) Kinetics of NR accumulation in DB-LP-GFP-NR-1 and DB-LP-GFP-NR-2 BCs, which have different affinities for NR, upon addition of 0.1 μM NR to the culture medium. D) BCs of the fully *de novo* protein containing a NR/NB-binding domain can be visualised by staining with NR or NB.

When expressed in HeLa cells and stained with NR or NB added to the cell media, DB-LP-GFP-NR-1 formed condensates that were dual coloured for both GFP and NR (Fig. 3A, row 1) or NB (Supplementary Fig. 13, row 1) binding. The parent protein without the binding site, DB-LP-GFP, did not show NR/NB fluorescence (Fig. 3A, row 2 and Supplementary Fig. 13, row 2). Furthermore, when imaged in real-time, addition of 0.1 μM NR to the cell medium resulted in rapid and efficient accumulation of NR in the DB-LP-GFP-NR-1 condensates (Fig. 3B). This is apparent from the increase in yellow in the merged images from the GFP and NR channels with the DB-LP-GFP-NR-1 construct (top row), compared with the separate green and red signals with the parent, DB-LP-GFP, construct (bottom row). This corresponded to a ≈65-fold increase in NR intensity over the control (Fig. 3C).

We compared the performance of DB-LP-GFP-NR-1 as a NR sponge to another variant, DB-LP-GFP-NR-2 (Supplementary Fig. 14), with a 10-fold tighter NR-binding site installed in the first LP domain (K_D,NR_ = 0.22 ± 0.04 μM^60^). This increased NR fluorescence to ≈150-fold over the control (Fig.4C). This capacity of the *de novo* BCs to act as small-molecule sponges and accumulate NR resulted in complete dominance of BCs over other cell structures (like lipid vesicles) that are typically stained by NR^61^, as observed for the reference parent sample of DB-LP-GFP (Fig. 3A and Supplementary Fig. 13, bottom rows).

**Fig. 4.**
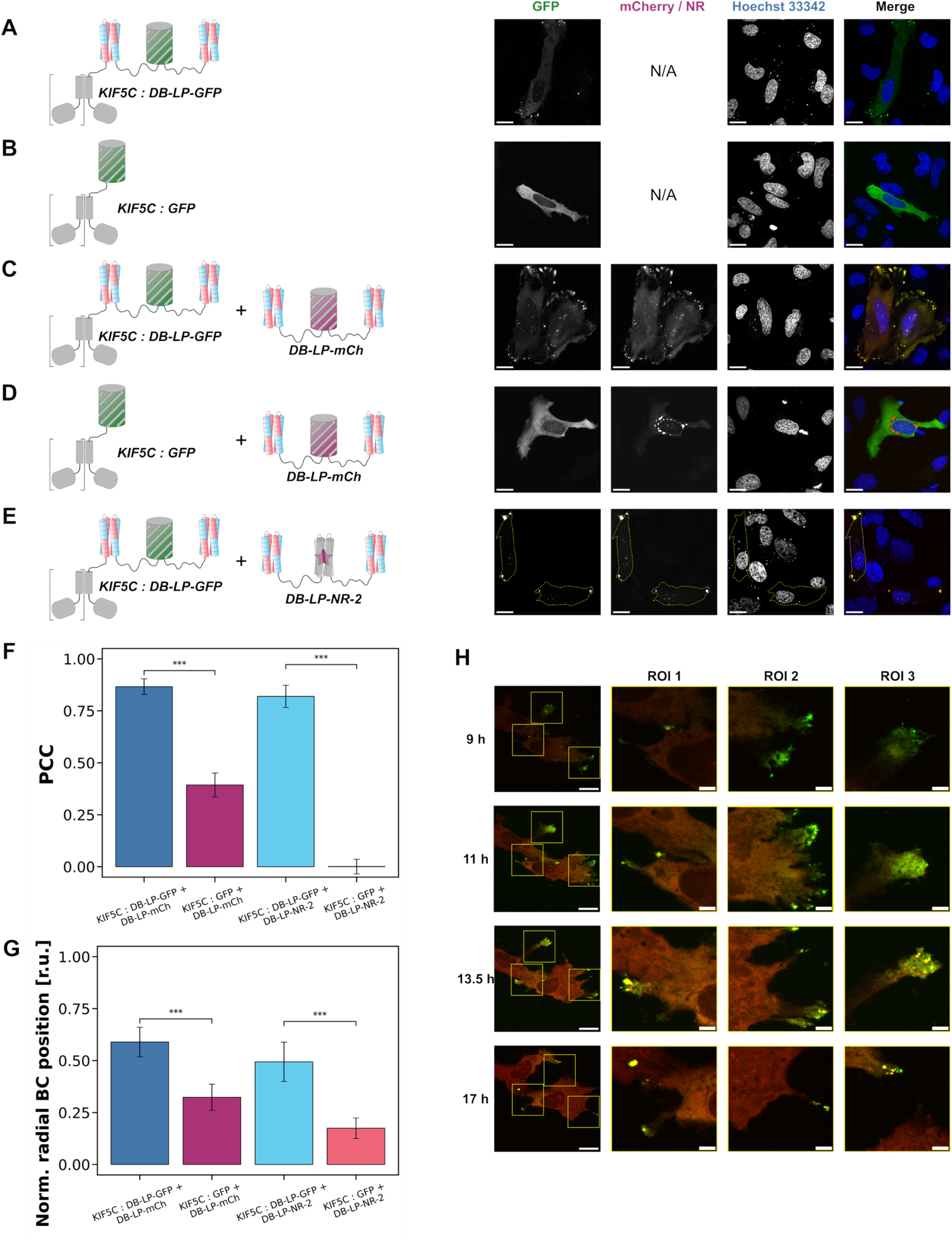
Differential localization of BCs using kinesin-1 motor proteins. Confocal fluorescence images of live HeLa cells expressing A) KIF5C:DB-LP-GFP; B) KIF5C:GFP; or co-expressing DB-LP-mCherry along with either C) KIF5C:DB-LP-GFP or D) KIF5C:GFP fusions to the kinesin motor protein in 24 hours after transfection. E) NR sponges can be localised at the cell periphery when the DB-LP-NR-2 is co-expressed with a motor-DB protein nucleating BC formation. F) Co-localization efficiency (expressed as Pearson correlation coefficient (PCC)) of the free component (75% by DNA), either DB-LP-mCherry (bars 1&2) or DB-LP-NR-2 construct (bars 3&4) in the red channel and the motor protein (25% by DNA) in the green channel either fused to a full length DB-LP-GFP (bars 1&3) or to a dummy cargo (GFP; bars 2&4). G) Quantification of the distribution of individual BCs between the nucleus (taken as 0) and the cell periphery (taken as 1). Bars correspond to BC formed by a free component (75% by DNA), either DB-LP-mCherry (bars 1&2) or DB-LP-NR-2 construct (bars 3&4) in the presence of an overexpressed motor (25% by DNA) either fused to a full-length DB-LP-GFP (bar 1&3) or to a dummy cargo (GFP; bars 2&4). H) Time-lapse imaging of HeLa Cells co-expressing KIF5C:DB-LP-GFP BC-nucleating protein and DB-LP-mCherry. Time after transfection is shown above the images. Scalebars = 20 μm, scalebars in images of selected regions of interest (ROI) = 5 μm.

We compared the time courses of NR accumulation in BCs with synthetic receptors with different affinities to the ligand, DB-LP-GFP-NR-1 and DB-LP-GFP-NR-2 (Fig. 3C). Interestingly, despite the >2-fold difference in overall uptake of NR by the two BCs, the effective half-time for NR accumulation was the same for both. This suggests that equilibration is limited by NR supply, and the sponge system can be described by a straightforward model where the observed on rate (*k*_obs_)— a compound of the rates of i) NR entry into the cell, ii) NR partitioning into BC, and iii) the *k*_on_ for binding—are similar regardless of the receptor, and the BC capacity for NR is determined by the binding constant, K_D,NR_.

Finally for these experiments, and to exploit the designed modular framework of the dumbbell system further and to utilise fluorophore-binding capacity fully, we removed the GFP reporter altogether and imaged the condensates directly. We replaced the GFP component of DB-LP-GFP with the NR binders to give DB-LP-NR-1 (Fig. 3D) and DB-LP-NR-2 (Supplementary Fig. 15). This used the NR-binding domains described above, but without surface charged patches for phase separation, and with overall charges matching GFP (*i.e.* -6 at pH 7.5 for both GFP and mCherry). When expressed in HeLa cells, these fully *de novo* dumbbell-like proteins formed condensates alone (Fig 3D) and when co-expressed with DB-LP-GFP (Supplementary Fig. 16A), with the former only visible when NR was added to the cell media.

### Dumbbell condensation can be controlled and localised in cells

Finally, we sought to control and localise the assembly of the DB-LP-GFP condensates in cells. To do this, we fused DB-LP-GFP to the *C* terminus of a motor protein, Fig. 4A ^13,62^; specifically, a truncated form of rat KIF5C kinesin (residues 1 – 405) containing only the motor domain and the neck coiled-coil region required for dimerization, which form a constitutively active motor that transports cargoes along microtubules to the cell periphery ^63^. When overexpressed in HeLa cells, the KIF5C(1-405):DB-LP-GFP fusion gave a phenotype of elongated cells expected for the active motor^64^, Fig. 4A. Importantly, condensates were only observed at the cell periphery for this fusion protein containing DB-LP-GFP and not with a dummy of KIF5C(1-405) fused to GFP alone, Fig. 4B.

To test for nucleation and growth of droplets by a free protein, we co-expressed the nucleating construct, KIF5C:DB-LP-GFP (25% by transfected DNA), with free DB-LP-mCherry (75% by DNA), Fig. 4C. In this case, the GFP and mCherry signals overlapped indicating that the motor-bound and free DB constructs co-localised with a Pearson’s correlation coefficient (PCC) close to 1 (Fig. 4F). Moreover, these dual-labelled puncta were mostly at the cell periphery compared to the control, KIF5C:GFP-only fusion, where the mCherry-containing BCs remained largely perinuclear (Fig. 4C&D). Both the relative position of individual BCs with respect to the cell periphery (Fig. 4G) and the distribution of DB-LP-mCherry fluorescence (Supplementary Fig. 17) supported these observations. Moreover, the process of BC nucleation and growth could be followed in real time (Fig. 4H). As shown on zoomed-in regions (ROI 1 – 3), at earlier time points, green KIF5C:DB-LP-GFP formed small puncta at the cell periphery that further co-localised with the red signal of DB-LP-mCherry and increased in size over time resulting in larger yellow BC.

Further, we substituted the free mCherry-containing dumbbell with the completely *de novo* designed constructs from the previous section; namely, DB-LP-NR-1 (Supplementary Fig. 16B-C) and DB-LP-NR-2 (Fig. 4E, Supplementary Fig. 16D). These were selectively recruited to BCs at the cell periphery by the KIF5C:DB-LP-GFP construct (Fig. 4G). Remarkably, the co-localization expressed as PCC increased from near-zero without the nucleator to almost complete with it (Fig. 4F). Thus, these experiments demonstrate that the *de novo* BCs can be assembled at a specific location in cells—in this case at the cell periphery exploiting a kinesin-based motor—where they can be used as small-molecule sponges.

## CONCLUSION

We have shown that properties of natural biomolecular condensates (BCs) can be captured in relatively straightforward *de novo* designed proteins built using understood, modular components. The modules can be substituted in predictable ways to alter properties of the proteins, and to bring functions to the BCs. Unlike many engineered, and the relatively few *de novo* designed phase-separating protein systems^4–9,11–17,37,65^—many of which rely on intrinsically disordered regions to drive phase separation^5,11,13,15,16,65^—the key building block for our BCs is a small *de novo* designed globular module. This brings predictability and designability to the system. Specifically, we use a *de novo* water-soluble four-helix bundle (4HB) built using clear sequence-to-structure relationships. This allows the surface residues to be altered rationally to dial in stickiness; that is to introduce weak, non-specific, protein-protein self-interactions. Whilst the resulting isolated domain remains fully soluble, joining two with a disordered linker gives dumbbell proteins that phase separate to form BCs *in vitro* and in cells. Detailed characterization of these BCs shows that they are reversible, highly dynamic, liquid-like droplets with physical properties very similar to those of natural BCs. Variants of the 4HB can be substituted in to add functions such as fluorophore binding. Effectively, this creates small-molecule sponges that sequester fluorophores allowing direct imaging in cells without any other labels. The location of the BCs can also be controlled by fusing a nucleating dumbbell to an engineered motor^13,62^, which transports the construct to the cell periphery where it recruits free dumbbell proteins to assemble hybrid BCs. In these respects, the *de novo* designed proteins and BCs add to the repertoire of phase-separating systems in cells^4,6–9,14,15,65^. Moreover, they provide a distinct means to disentangle sequence-to-structure/function relationships underlying the phenomenon. Furthermore, they present new possibilities to develop compartmentalization strategies to manipulate subcellular processes in cell biology^7,13,15,66,67^, and to introduce synthetic processes to augment endogenous cell functions for engineering biology^7,13,15,62,65^.

## METHODS

### General

All solvents, chemicals, and reagents were purchased from commercial sources and used without further purification. Luria-Broth (LB), antibiotics, IPTG were purchased from Thermo Fisher. All other chemicals were reagent grade and purchased from Sigma-Aldrich. Protein biophysical characterization (unless specified otherwise) were recorded in a 50 mM Tris-HCl, 500 mM NaCl, pH 7.5.

### Computational tools

The code to run the RASSCCoL protocol can be found in the Woolfson group GitHub (https://github.com/woolfson-group/RASSCCoL_no_RF/tree/master). Because the scaffold used to design NR binders^60^ and patchy sc-apCC-4 variants was the same, to generate the patchy binders combining both features we mutated the core of the patchy sc-apCC-4-LP design to match the one from either sc-apCC-4-NR-1 or sc-apCC-4-SN38-1^60^

To design NR binders with the net charge of -6 matching GFP or mCherry, we used the original designs sc-apCC-4-NR-1 and sc-apCC-4-SN38-1 ^60^ and changed some of the f-positions in the heptads containing lysines in *b-* and *c-*positions to glutamates and thus ensuring well-mixed charged residues on the protein surface.

AlphaFold2^68,69^ (ColabFold^70^ v1.5.5) using single-sequence mode (for sc-apCC-4 variants) or with MSA (for designs containing natural proteins, e.g. GFP), and 20 recycle steps was used to generate models for protein designs. The top rank model was relaxed using Amber. AlphaFold3^71^ was accessed via the web server https://alphafoldserver.com with default settings.

PROPKA 3^72–75^ (v3.5.0) obtained from GitHub (https://github.com/jensengroup/propka-3.0) was used to assign protonation states at various pH. The Docker version of MaSIF ^76^ available on GitHub (https://github.com/LPDI-EPFL/masif/tree/master) was used to predict chemical and geometric fingerprints of patchy designs. APBS^77^ calculations were run using the web server (https://server.poissonboltzmann.org/). APBS is also run as a part of MaSIF pipeline (Extended Data Fig. 1).

### Transfection and imaging of HeLa and HEK 293T cells

The linear DNA fragments for the designer constructs were synthesised by GeneArt or IDT and then subcloned in pTwist CMV vector (Twist Bioscience) for expression in HeLa cells. When required, PCR reactions were carried out using Q5 High Fidelity Hot Start DNA Polymerase (New England Biolabs (NEB)), following manufacturers’ instructions. PCR products were purified using Monarch® PCR & DNA Cleanup Kit (NEB). Restriction digest reactions were carried out using NEB restriction enzymes at 37 °C for 1 h. Ligation reactions were carried out using T4 DNA Ligase and Rapid ligation buffer (Promega) for 1-2 h at 16 °C. Transformation was carried out in competent (subcloning efficiency) *Escherichia coli* cells (NEB® 5-alpha from NEB or DH5α from Invitrogen) for 30 min on ice, followed by heat shocking (30 s, 42 °C), and a further 5 min on ice. Cells were recovered by incubating in LB media at 37 °C for 1 h and then plated on LB-agar plates containing the ampicillin.

Plasmid DNA samples for transfection were purified using Plasmid Midi Kit according to the manufacturer’s instructions (QIAGEN) and verified by DNA sequencing using Rapid Sequencing Service provided by Source Bioscience.

HeLa cells were sourced from ECACC (UK Health Security Agency) and HEK293T cells were a gift from Professor Pete Cullen (University of Bristol) sourced from the American Type Culture Collection (ATCC). Both cell lines were maintained in high glucose Dulbecco’s Modified Eagle’s Medium with 10% (v/v) foetal calf serum (Sigma-Aldrich) and 5% penicillin/streptomycin (PAA) (herein referred to as DMEM) without phenol red at 37 °C and 5% CO_2_. For transfection, cells were seeded on tissue culture treated 35 mm CELLview cell culture dishes with 10 mm glass bottom (Greiner) at a density of 1×10^5^ cells per well and incubated at 37 °C and 5% CO_2_ for 16 h prior to transfection. Cells were transfected with 0.8 μg of the indicated plasmid DNA using Effectene transfection reagent according to the manufacturer’s instructions (QIAGEN). After transfection cells were incubated at 37 °C, 5% CO_2_ for 6 h, washed with fresh DMEM and incubated further for 18 h. Before imaging, cells were washed with PBS and, when specified, stained with Hoechst 33342 (H3570, 1 μg/mL), LysoTracker Deep Red (L12492, 0.05 μM), MitoTracker Red FM (M22425, 0.1 μM), CellMask Deep Red (A57245, 1X), ER-Tracker Blue-White DPX (E12353, 0.1 μM) (all from Thermo Fisher Scientific), and Nile Red (reagent grade, Sigma-Aldrich, 0.1 μM) at 37 °C for 30 minutes, washed again, and then the medium was replaced for fresh DMEM.

Confocal images were collected using Olympus IXplore SpinSR system with a 60× objective lens or Leica SP5II microscope with a 63× objective lens at 37 °C using the following imaging channels (λex-max λem/width λem): 405-447/60, 488-525/50, and 561-617/73 nm. Figures were assembled using the Fiji distribution of ImageJ2^78^.

### Image analysis

Signal distribution was quantified using a custom workflow for the ModularImageAnalysis (MIA) plugin for ImageJ^78–80^. The analysis was performed on z-projected (maximum) images which were first processed with a rolling ball filter (radius = 200 px) to remove background signal. Cells were segmented using a two-step approach using both the nuclear and free component channels. For the initial nuclear detection the image was intensity normalised and passed through a 2D median filter (radius = 2 px) before candidate nuclei were detected using Cellpose-SAM^81^. Any objects smaller than 50 μm^2^ were excluded from further analysis and the remaining nuclei were optionally manually edited to correct any segmentation errors. Cells were likewise detected using Cellpose-SAM on images which had previously been intensity normalised and converted to a log scale. In this instance, any cells smaller than 250 μm^2^ or with intensity below a user-defined threshold were excluded from further analysis. The remaining cells were also subject to optional manual editing.

Detected nuclei and cells were associated based on spatial overlap. Each cell was subdivided into a series of concentric bands^82^, starting at the nuclear surface and having 2 μm width. The intensity of the BC channel (both with and without background subtraction applied) was measured for each band. Additionally, individual BC objects were segmented using an intensity thresholding-based approach whereby the BC channel was 2D median filtered (radius = 2 px) and binarised using the Otsu auto-threshold algorithm ^83^. BC objects were detected using connected components labelling ^82^ and subject to a minimum 1 μm^2^ size threshold. The fractional distance between the nuclear surface and cell surface (both approximated as convex hulls) was calculated for each BC object.

### Protein purification

Proteins were expressed and purified from *E. coli* BL21 Star^™^ (DE3) cells transformed with pET-28a(+) plasmids containing the respective genes. For protein purification, 1 to 12 L of LB was inoculated 1:100 from an overnight culture, induced with 0.4 mM IPTG at OD 0.6, and then grown at 18 °C, shaking at 200 rpm. Cell pellets were resuspended in buffer containing 50 mM Tris-HCl pH 7.5, 500 mM NaCl, 25 mM imidazole, 1 tablet of cOmplete protease inhibitor (Roche), and lysed by sonication on ice (3 s on, 3 s off, 70% amplitude, 10 minutes). The lysate was centrifuged (18000 xg, 45 minutes) and the supernatant filtered through a 0.45 μM filter to clarify. Protein purification was performed using an Äkta Pure (Cytiva) at 4 °C, with chromatograms monitored at 280 nm. The clarified lysate was applied to a HisTrap HP (Cytiva) immobilised metal affinity chromatography (IMAC) column, pre-equilibrated in 50 mM Tris-HCl pH 7.5, 500 mM NaCl, 25 mM imidazole. The column was washed until A_280_ was re-stabilised (typically 3 – 4x the column volume), before eluting the bound protein up-flow with imidazole (50 mM Tris-HCl pH 7.5, 500 mM NaCl, 500 mM imidazole). The combined protein fraction was desalted using a HiPrep 26/10 desalting column (Cytiva) into 50 mM Tris-HCl pH 7.5, 500 mM NaCl and then treated with recombinant TEV protease to remove the N-terminal His-tag. Cleavage by TEV protease was performed at room temperature overnight (for sc-apCC4-SP and sc-apCC4-LP) or at 4 °C over 16-40 hours (for DB-SP-GFP and DB-LP-GFP).

The cleaved protein was purified by application to a HisTrap HP column and collection of the flow-through. Cleavage was confirmed by SDS-PAGE and staining using Coomassie-blue. Recombinant proteins were further purified by size-exclusion chromatography using a HiLoad 16/600 Superdex 75pg (for sc-apCC4-SP and sc-apCC4-LP) or 200 pg (for DB-SP-GFP and DB-LP-GFP) exclusion column (Cytiva) with a flow rate of 1 ml/min. Size-exclusion was performed using a 50 mM Tris-HCl pH 7.5, 500 mM NaCl running buffer and elution monitored by A_280_. Protein fractions were identified by SDS-PAGE and the relevant fractions pooled. For characterisation, the proteins were used fresh after purification.

### TEV protease purification

TEV protease was expressed and purified from E. coli BL21 Star^™^ (DE3) cells transformed with pDICa-MBP-His-TEV (AmpR). Cultures were grown in LB medium supplemented with ampicillin at 37 °C to an OD600 of 0.6, induced with 0.4 mM IPTG, and incubated overnight at 37 °C. Cells were harvested by centrifugation (4000 rpm, 20 min, 4 °C), resuspended in binding buffer (50 mM phosphate pH 8.0, 300 mM NaCl, 10 mM imidazole, 5% glycerol), and lysed by sonication. Clarified lysate (12,000 rpm, 20 min, 4 °C) was subjected to immobilised metal affinity chromatography (IMAC) on an ÄKTA system, and TEV protease was eluted with buffer containing 300 mM imidazole. Fractions containing TEV were pooled and further purified by size-exclusion chromatography (Superdex 200 pg 16/600) in storage buffer (25 mM Tris-HCl pH 7.6, 100 mM NaCl, 10% glycerol). TEV protease eluted at 85–95 ml (peak ∼88 ml) and was concentrated to ∼2 mg/ml (A280 ≈ 2.0). Purified protein was aliquoted and stored at −80 °C.

### Circular-dichroism spectroscopy

Circular-dichroism (CD) data were collected on a JASCO J-810 spectropolarimeter fitted with a Peltier temperature controller (Jasco UK). Full spectra were measured between 190 and 260 nm with a 1 nm step size, 100 nm·min^−1^ scanning speed, 1 nm bandwidth and 1 second response time. Spectra were measured at 5 °C unless otherwise stated. Baselines recorded using the same buffer, cuvette and parameters were subtracted from each dataset. The spectra were converted from ellipticities (deg) to mean residue ellipticities (MRE, (deg.cm^2^.dmol^−1^.res^−1^)) by normalizing for concentration of peptide bonds and the cell path length using the equation:

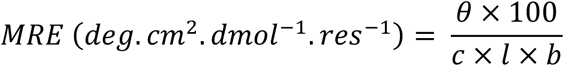

Where the variable *θ* is the measured difference in absorbed circularly polarised light in millidegrees, *c* is the millimolar concentration of the specimen, *l* is the path-length of the cuvette in cm and *b* is the number of amide bonds in the polypeptide, for which the *N*-terminal acetyl bond was included but not the *C*-terminal amide. Protein concentration was determined at 280 nm (ε_280_(Trp) = 5690 cm^−1^, ε_280_(Tyr) = 1280 cm^−1^)^84^ using a Nanodrop 2000 (Thermo) spectrometer.

### Small-angle X-ray scattering (SAXS) and Size-exclusion chromatography small-angle X-ray scattering (SEC-SAXS)

SAXS data for sc-apCC-4-SP and sc-apCC-4-LP were obtained at European Synchrotron Radiation Facility (ESRF, Grenoble, France) on beamline BM26. Samples were prepared to 1 mg/mL in 50 mM Tris-HCl pH 7.5, 500 mM NaCl. SEC-SAXS data for DB-SP-GFP and DB-LP-GFP were obtained at Diamond Light Source on beamline B21. Samples were prepared to 5 mg/mL in 50 mM Tris-HCl pH 7.5, 500 mM NaCl. A Shodex KW-402.5 column was equilibrated in the same buffer at 4 °C.

For both SAXS and SEC-SAXS, buffer subtraction and data merging was performed with ATSAS. qmin was taken as the first point of the linear Guinier region, qmax was calculated using ShaNum through ATSAS interface^85^. FoXS^86,87^ software (Sali Lab) was used for fitting experimental scattering profiles to AlphaFold2 models (Table S2) of sc-apCC-4-SP and sc-apCC-4-LP. MultiFoXS^86,87^ software (Sali Lab) using a monomer model was used for fitting experimental scattering profiles to AlphaFold2 models (Table S2) with flexible residues in the linker regions of the dumbbell proteins (10,000 conformations). The goodness of fit was described by χ^2^.

### Analytical Size-exclusion chromatography (SEC)

Analysis of protein by SEC chromatography was performed using Jasco HPLC instrument equipped with a Sephadex 75pg Increase column and UV-1575 UV-detector at the flow rate of 0.8 mL/min in 50 mM Tris pH 7.5, 500 mM NaCl as an eluent. Protein samples at 1-2 mg/mL were prepared in the same eluent solution. The UV traces were recorded at the wavelength of 230 nm.

### Analytical ultracentrifugation (AUC)

AUC was performed on a Beckman Optima X-LA or X-LI analytical ultracentrifuge with an An-50-Ti or An60-Ti rotor (Beckman-Coulter). Buffer densities, viscosities and protein partial specific volumes (v̅) were calculated using SEDNTERP (http://rasmb.org/sednterp/). For sedimentation velocity (SV), protein samples were prepared at 80 μM sc-apCC-4-SP and sc-apCC-4-LP and 15 μM DB-LP-GFP concentration and placed in a sedimentation velocity cell with 2-channel centrepiece and quartz windows. The samples of sc-apCC-4-SP and sc-apCC-4-LP were centrifuged at 50 krpm and DB-LP-GFP – at 40 krpm, at 20 °C, with absorbance scans taken over a radial range of 5.8 – 7.3 cm at 5 min intervals to a total of 120 scans. Data from a single run were fitted to a continuous c(s) distribution model using SEDFIT^88^ at 95% confidence level. Residuals for sedimentation velocity experiments are shown as a bitmap in which the grayscale shade indicates the difference between the fit and raw data (residuals < -0.05 black, > 0.05 white). Good fits are uniformly grey without major dark or light streaks. Sedimentation equilibrium (SE) experiments were performed at the same protein concentrations as SV experiments in 110 μL at 20 °C. The experiment was run in triplicate in a six-channel centrepiece. The samples were centrifuged at speeds in the range of or 16 – 48 krpm and scans at each recorded speed were duplicated after equilibration for 8 hours. Data were fitted using SEDPHAT^89^ to a single species model. Monte Carlo analysis was performed to give 95% confidence limits.

### Dynamic light scattering

For DLS measurements, the proteins were purified as mentioned previously and further characterised in the eluent buffer to match exactly the composition. Buffers were filtered through Anatop 0.02 μm filters (Whatman) were used for preparation of different protein concentrations. On the day of the experiment, the proteins were concentrated to 15 – 30 mg/ml concentration using Amicon Ultra Centrifugal filters (Merck) via short (2 – 5 min) cycles at the speed ≤ 3000 xg at 20 °C, and then centrifuged for 60 – 90 minutes at 17,000 ×g at room temperature to remove any pre-formed aggregates in solution.

An ALV/CGS-3 goniometer with a HeNe laser operating at a wavelength of 632.8 nm, an optical fibre based detector and an ALV/LSE-5004 Light Scattering Electronics and Multiple Tau Digital Correlator were used for DLS measurements. The temperature was kept constant at 20 °C during data acquisition using a Thermo Scientific DC30-K20 water bath connected to the instrument and measured with a Pt-100 probe immersed into the index matching fluid vat. DLS measurements were carried out for 30 – 60 minutes at a scattering angle of 90 °C at each protein concentration. The protein concentration was determined for the sample after the last measurement using Cary-100 (Agilent) UV-Vis spectrometer based on the extinction coefficients at 280 nm (ε_280_(Trp) = 5690 cm^−1^, ε_280_(Tyr) = 1280 cm^−1^)^84^.

Volume fraction is calculated using the expression *c* = *ϕ*×*n* where *c* is the concentration in mg ml^−1^, *ϕ* is the volume fraction and *n* is the partial specific volume equal to 7.555 10^−4^ for sc-apCC4-SP and sc-apCC4-LP, 7.421 10^−4^ ml·mg^−1^ for DB-SP-GFP and DB-LP-GFP, and 7.325 10^−4^ ml·mg^−1^ for GFP, respectively, as calculated using sedfit software.^88^

To calculate the hydrodynamic radius from the diffusion data, the Stokes–Einstein–Sutherland equation for spherical particles was used:

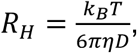

Where *k*_B_ is the Boltzmann constant, *T* is the absolute temperature, *D* is the diffusion coefficient, and *η* is the dynamic viscosity.

### Fluorescence recovery after photobleaching

Fluorescence recovery after photobleaching (FRAP) of GFP was performed using a Leica SP8 AOBS confocal with a 65 mW Ar laser exciting at 488 nm at 22 °C. For each bleaching measurement 10-20 images were taken before bleaching and the mean intensity recorded as the pre-bleach fluorescence intensity. Bleaching was performed using a 1 ms laser burst at 50-100% laser power, followed by imaging every 76 ms for 30 – 60 s to record fluorescence recovery. Data analysis was performed in Python. The fluorescence intensity of the background was subtracted from all measurements. For each bleaching measurement, recovery was normalised relative to the mean fluorescence intensity before bleaching (normalised to 1), and the minimum fluorescence intensity measured immediately after bleaching (normalised to 0) to allow comparison between different bleaching experiments. Bleaching was further normalised to a reference droplet to compensate for bleaching during measurement. For *in vitro* measurements, de-mixed droplets were placed on a clean glass slide and covered with a cover slip before imaging. *In vitro* conditions were 4 mg/ml DB-LP-GFP, 150 mM NaCl, 50 mM Tris-HCl pH 7.5, and 2% PEG 3350. For in-cell measurements, FRAP was performed on live HeLa cells prepared in the same way as for imaging.

The raw FRAP data from bleached ROI1 after background subtraction was normalised to 100% using the average pre-bleach intensity in ROI1 and to 0% using the first point after bleaching. Bleaching in the sample during measurement was taken into account by normalisation to the background-subtracted relative signal in a non-bleached ROI. The processed data was then fitted to mono-, bi- and stretched exponential models with respective recovery half-time *τ*_1/2_:

| Model | Function | Half-time $\tau_{1/2}$ [s] |
| --- | --- | --- |
| Monoeexponential | $I(t) = a(1 - e^{-t/b}) + c$ | $b \times \ln(2)$ |
| Biexponential | $I(t) = a_1(1 - e^{-t/b_1}) + a_2(1 - e^{-t/b_2}) + c$ | $f(\infty) + 0.5[f(0) - f(\infty)]$<br>(solved numerically) |
| Stretched exponential<br>(Anomalous diffusion) | $I(t) = a(1 - e^{-t/b}) + c$ | $b \times [\ln(2)]^{1/\alpha}$ |

### FCS measurement in live cells

FCS measurements were performed in Leica SP8 inverted epifluorescence microscope equipped with pico quanta autocorrelator. A water immersion objective lens (63x/1.3) and white light laser (adjusted the excitation wavelength and detection window based on the FCS probe fluorophore excitation/emission wavelength) used for collecting the photons.

Although the protein condensate was co-expressed with low level of DB-LP-mCherry, the amount of fluorophore was excess for FCS measurement. Therefore, before each measurement some amount of mCherry was bleached using high laser intensity. The amount of bleaching was decided by looking the count rate with relatively lower laser power (in the settings only used 2% of laser power). To reduce the artifact the larger visible BCs inside HeLa cell were chosen to measure FCS. The excitation volume was positioned inside the condensate based on the confocal images from DB-LP-GFP. Once the FCS volume is placed inside the condensate, measurements recorded for 10 s from multiple points within the condensate. Five repeated measurements taken from each point. From the fluorescence intensity fluctuations data, the autocorrelation curve generated using a python package based on^90^. Even though lower laser power used, some amount of photo bleaching observed during the data acquisition. Before calculating the autocorrelation, the fluorescence intensity fluctuations data were treated for photobleaching correction^91^. The excitation volume was calibrated using Rh6G dye in water. The normalised autocorrelation curves measured from at least 5 cells were averaged before performing the fitting.

### Fitting the data

Anomalous diffusion model used to fit the data: 1-component and 2-component diffusion model^92^.

1- component anomalous diffusion model:

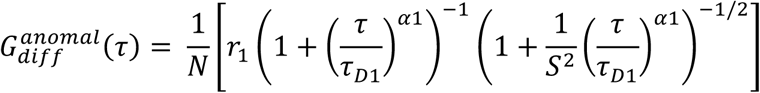

2- component anomalous diffusion model:

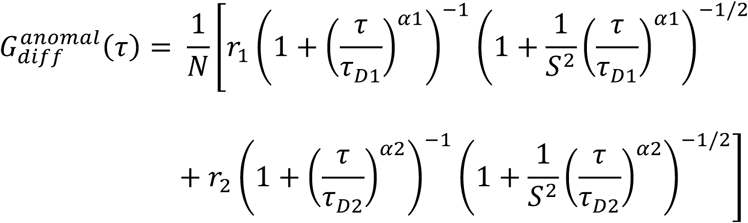

Where N is the average number of probe dye molecules in the excitation volume, *τ_D_*_1_ and *τ_D_*_2_are the diffusion time of slow and fast moving species, *r*_1_ and *r*_!_are the mole fraction of slow and fast moving species. *⍺*1 and *⍺*2 are anomalous factors. S is the structure factor of the system which estimate from the detection volume. After the FCS curve fitting the diffusion coefficient can be estimated using the equation

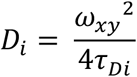

Where *ω*_56_ is the lateral radius of detection volume estimate from the diffusion of Rh6G dye in water.

### Estimated hydrodynamic radius

For globular proteins, the ratio of *R*_g_ and *R*_h_ is generally found to be 0.775^93^. Rg was computed in Pymol software using Psico library. For example, for mCherry *R*_g_ was calculated to be 1.65 nm and hence *R*_h_ is 2.13 nm.

## Data availability

Data from this study are openly available in Zenodo at DOI: 10.5281/zenodo.17204472.

## Acknowledgments

A.V.R. and R.K.T.P. were supported by a Leverhulme Trust grant to J.J.M. and D.N.W. (RGP-2021-049). R.P. is supported by a BBSRC-funded PhD studentship and by Rosa Biotech through the South West Biosciences Doctoral Training Partnership (BB/T008741/1). K.O. is supported by a BBSRC grant to Prof. Nigel Scrutton and D.N.W. (BB/X003027/1).J.J.C. was supported by an Engineering and Physical Sciences Research Council (EPSRC) program grant to Prof. Graham Leggett and D.N.W. (EP/T012455/1).

We thank Wolfson Bioimaging Facility staff for their assistance with the cell imaging; Dr Emily Sakamoto-Rablah for assistance with DLS measurements; Prof. Mark Dodding and Dr Jessica Cross for the kinesin constructs; and members of the Woolfson and McManus groups for many valuable discussions.

## Authors contributions

D.N.W., J.J.M, and A.V.R. conceived the project. A.V.R. and D.N.W designed the proteins. A.V.R. produced and characterised the proteins *in vitro* and in cells. R.K.T.P. performed FCS and condition screen imaging. K.O., R.P., J.J.C, and A.V.R designed the chromophore-binding proteins. S.J.C. developed programming code for automated image analysis. A.V.R., J.J.M. and D.N.W. interpreted the data and wrote the paper, and all authors contributed to its revision. J.J.M. and D.N.W. raised the funding.

## Additional information

Extended Data Figs. 1-3, Supplementary Tables 1–6, Supplementary Figs. 1–17, Supplementary Video 1, and references

## Extended data

**Extended Data Fig. 1.**
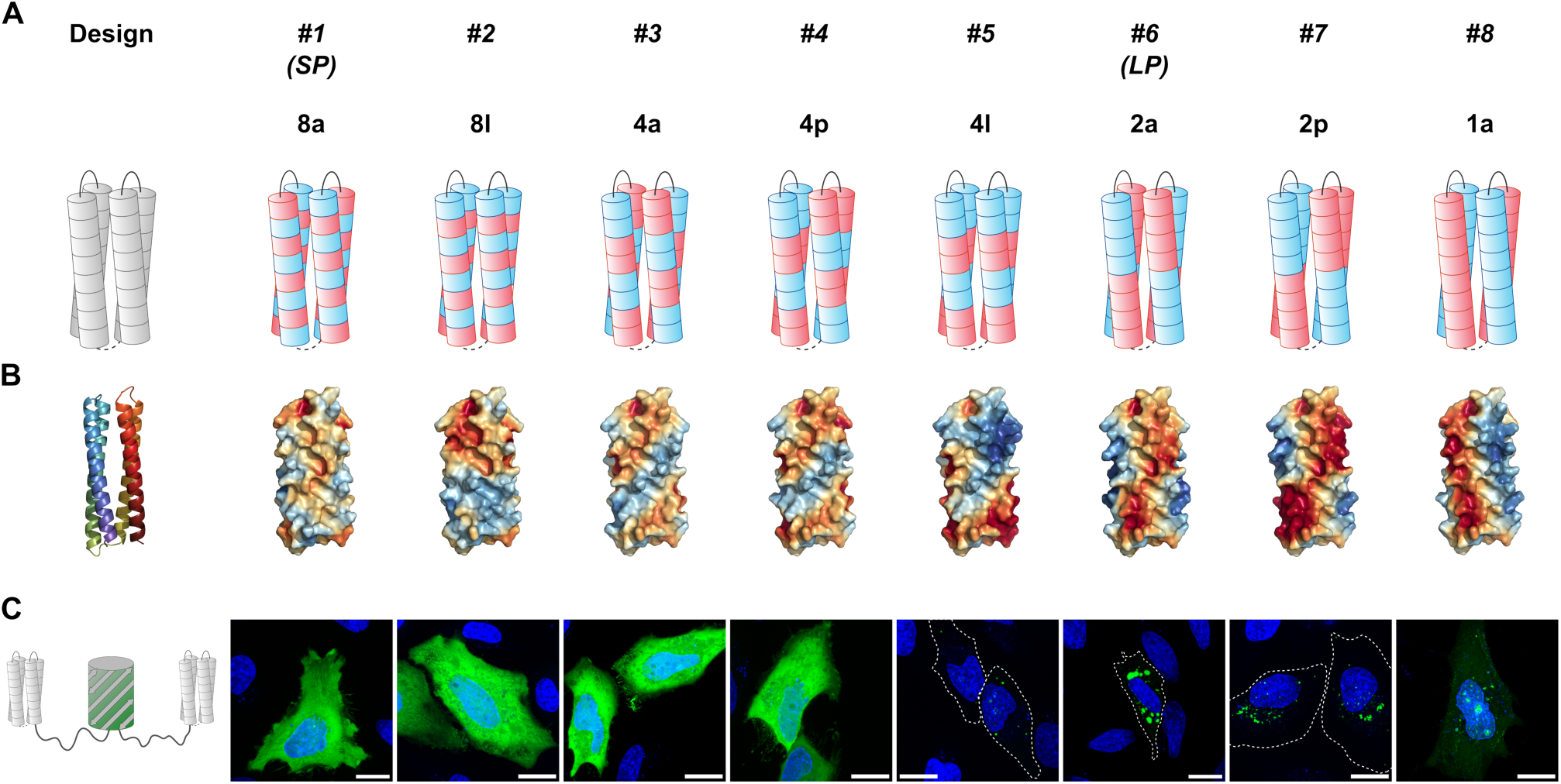
Rational design of sc-apCC-4 variants with increasing size of charged patches on their surfaces using *b*, *c*, and *f* positions of heptad repeats. (A) Schematics showing one *f-* or both neighbouring *b-* and *c-* positions as single slices of the helices represented by cylinders. This way, each helix has eight slice (either *f* or *bc*) that can be occupied either by Lys (K) or Glu (E): (*f*)(*bc*)(*f*)(*bc*)(Q@*f*)(*bc*)(*f*)(*bc*)(*f*), with the middle one always taken by a glutamine residue (Q@*f*). The systematic names start with a digit that denotes the number of a smallest repeat unit of either E or K type along the helix. This is followed by a letter describing the arrangement of the repeat units within each slice: alternating – a, layered – l, paired – p. (B) Surface potential maps calculated using APBS^94^ for all variants. (C) Confocal images of live HeLa cells transiently expressing the dumbbell proteins containing two copies of the 4-helix bundles shown in (A) and (B).

**Extended Data Fig. 2.**
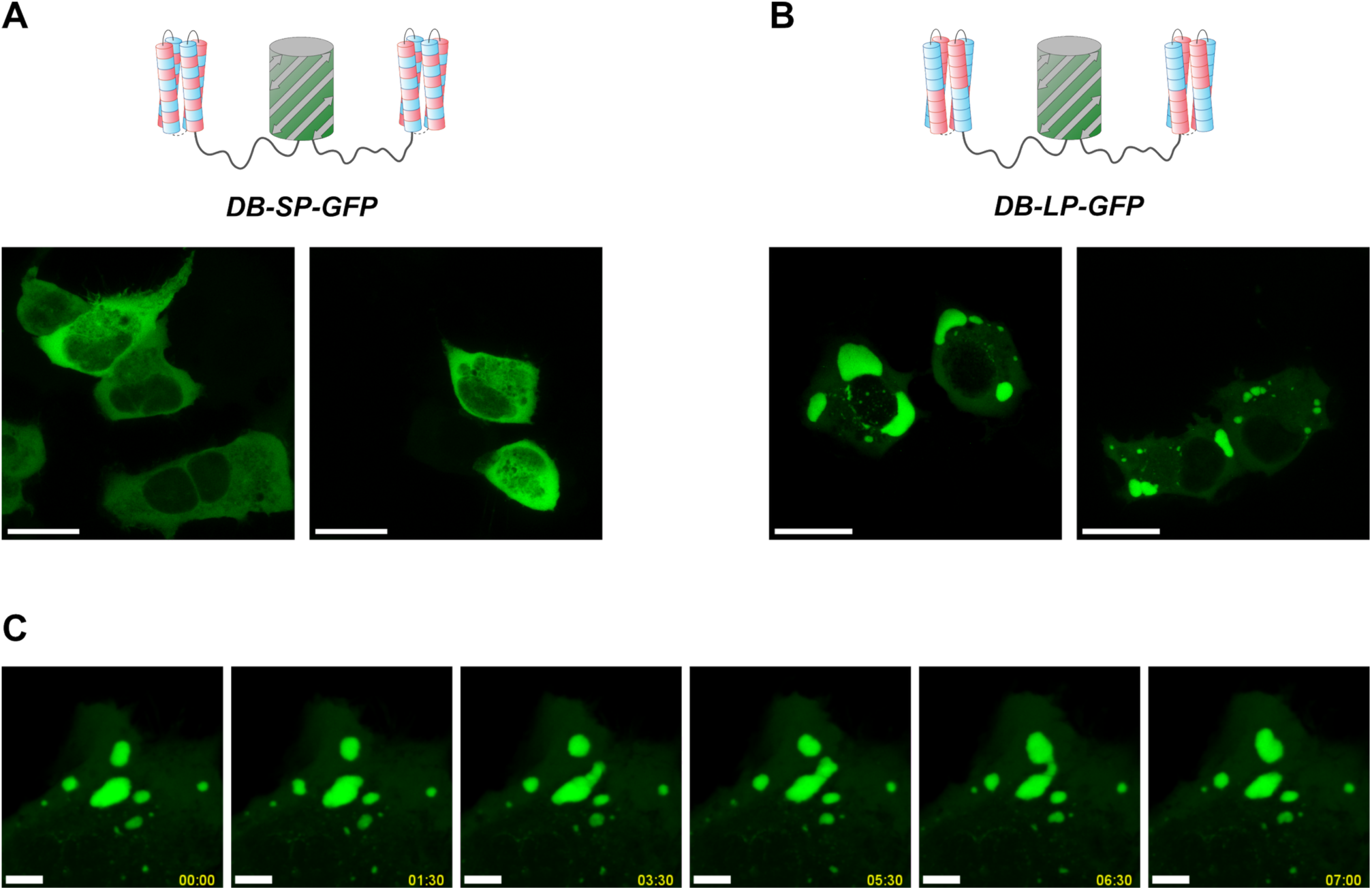
(A-B) Confocal images of HEK cells that transiently overexpressed the designer proteins. A) DB-SP-GFP containing sc-apCC-4-SP with small charge patches shows diffused fluorescence whereas B) DB-LP-GFP with large patches on sc-apCC-4-LP forms BCs. Scalebar = 20 μm. (C) Time-lapse imaging showing coalescence, fission, and flowing of BC formed by DB-LP-GFP in HeLa cells. Time format mm:ss. Scalebar = 5 μm. For the movie from live-cell imaging see Supplementary Video S1.

**Extended Data Fig. 3.**
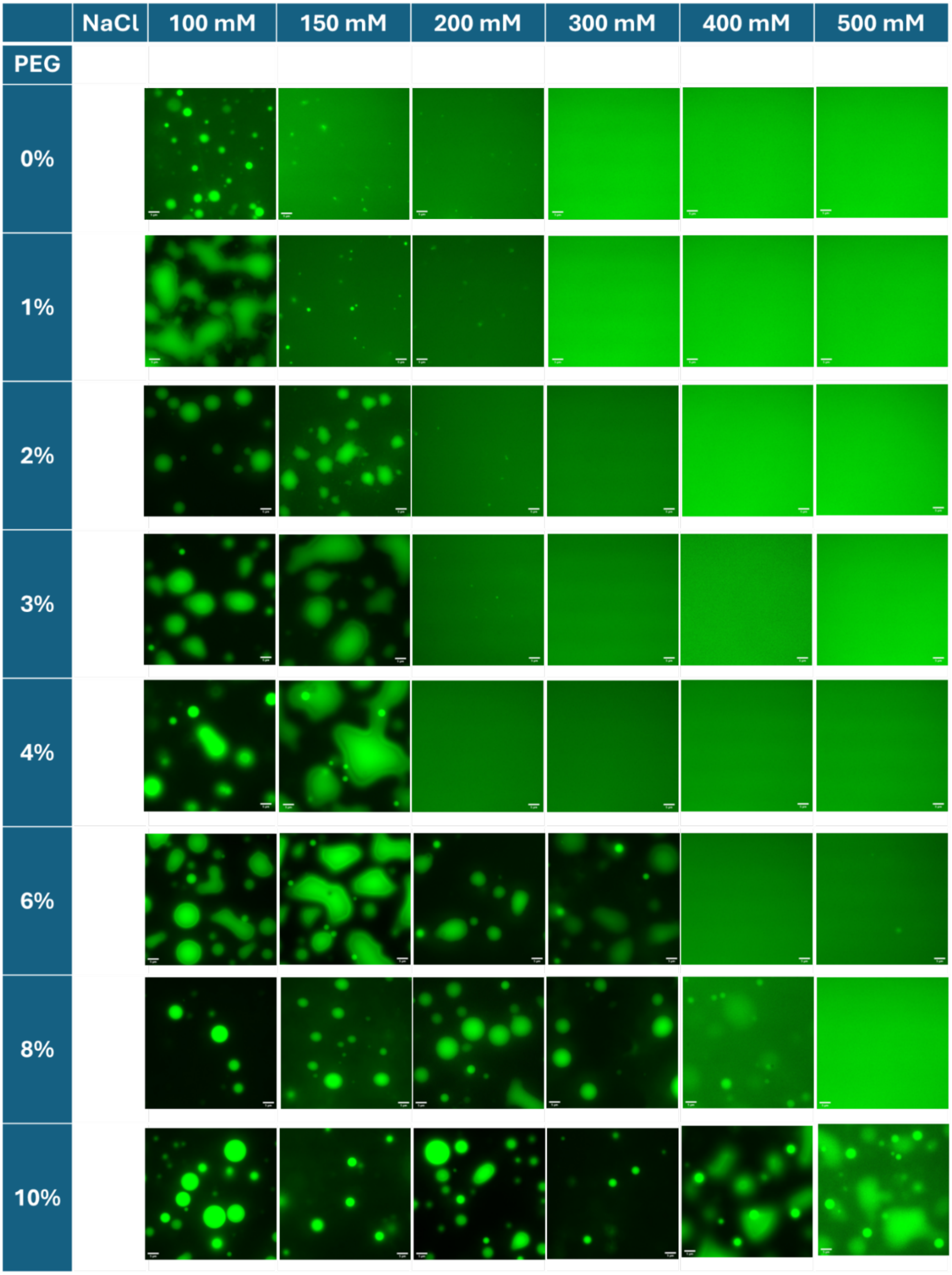
Fluorescence images of DB-LP-GFP solutions at various NaCl (100 – 500mM) and PEG 3350 (0 – 10%) concentrations. DB-LP-GFP concentration (2.5 mg/mL) and 50 mM Tris buffer at pH 7.5 were fixed. Scale bar 5 μm. RT.

